# Local activation of Cxcl12a signaling controls olfactory placode morphogenesis in zebrafish embryos

**DOI:** 10.1101/2024.12.20.629493

**Authors:** Marie Zilliox, Gaëlle Letort, David Sanchez, Christian Rouviere, Pascale Dufourcq, Frédérique Gaits-Iacovoni, Anne Pizzoccaro, Violaine Roussier-Michon, Patrick Blader, Julie Batut

## Abstract

Morphogenesis and cell-type differentiation are highly coordinated in sensory organs to ensure their function. Morphogenesis of the olfactory epithelium (OE) in zebrafish provides a unique model to study this process as undifferentiated cells aligned around the anterior neural plate mature into clusters of early olfactory neurons across a short time scale. While the Cxcl12a/Cxcr4b signaling pathway drives this process, what constraints on pathway activation apply during morphogenesis are unclear. We developed a mathematical model recapitulating Cxcl12a-mediated OE morphogenesis. Restoring Cxcl12a expression in mutants for the ligand rescues correct morphogenesis both in silico and in vivo. However, where expression of the ligand is restored is crucial for rescue, a point not predicted by our model and suggesting an unexpected level of pathway activation control. Analysis of a Cxcr4b activation reporter supports this idea. We concluded that mosaic and heterochronic Cxcl12a activation along the anteroposterior axis sculpts the olfactory epithelium.

## Introduction

Sensory organs of the vertebrate head permit the detection of an individual’s environment. These specialized structures develop early during embryogenesis and are composed largely of neurons and support cells derived from so-called placodal ectoderm and cranial neural crest ^1^. Once formed, each organ displays a specific morphology that is precisely adapted to their distinct function. Each organ is composed of specific sensory cell-types that are also tailored to their function, with olfactory neurons being found in the olfactory epithelium of the nose and sensory hair cells in the inner ear, for instance. Defects in either the morphogenesis of these organs or the correct specification of their sensory cell types can lead to sensory defects such as anosmia or deafness ^2,3^.

The zebrafish olfactory epithelium provides a highly tractable system to study the coordinated development of organ form and cell-type diversity. During zebrafish embryo development, olfactory placode precursors are located at the neural plate border as early as 12 hours post-fertilization (hpf), and form a horseshoe around the telencephalon ^1,4,5^. This neural border will give rise to the cephalic neural crests (CNCs) and cranial placodes. Cells from the olfactory territory will then assemble into the olfactory placode by actively converging and forming two tight clusters on either side of the anterior neural tube between 18 and 21 hpf ^6^. Then, a passive lateral movement will give its shape to the cluster starting from 20 hpf ^7^. Concomitant to cell convergence, two successive waves of neurogenesis exist, together with three signaling pathways, Cxcl12a/Cxcr4b, Slit/Robo and Netrin/DCC, that are hierarchically required for axonal guidance of OSNs ^4^. However, the process of placode formation between 12 and 24 hpf remains poorly characterized. Previous studies have shown that disruption of chemokine signaling through mutations of the chemokine receptor gene *cxcr4b* (*cxcr4b^t26035-/-^)* or its ligand *cxcl12a* (*cxcl12a^t30516-/-^)* lead to defects in the assembly of the olfactory placodes, an early step in the development of the olfactory epithelium ^8^. Our previous work indicates that two bHLH proneural transcription factors, Neurog1 and Neurod4, share a redundant role in the birth of Early Olfactory Neurons and the Olfactory Sensory Neurons during the same developmental time window ^9^. Recently, we published that Cxcr4b is directly regulated by Neurog1 ^10^. As such, Neurog1 coordinates both morphogenesis and neuronal specification in this system. We know from previous work that *cxcr4b* is expressed by EONs arranged along the horseshoe, while its ligand, *cxcl12a*, is expressed in the adjacent anterior neural plate/tube ^8^. How EONs perceive this signal and what parameters control the morphogenesis of the olfactory epithelium are still under debate.

To explore how chemokine signaling controls the convergence of cells to form olfactory placodes further, we combined mathematical modeling with *in vivo* experiments; the ability to employ mathematical simulations has become a powerful tool to select and test likely hypotheses concerning developmental mechanisms hidden in the complexity of *in vivo* systems ^11^. Using a simple theoretical framework, we tested different distributions of a source of chemotaxis, mirroring the activity of the Cxcl12a/Cxcr4b pathway, and followed if/how they affect the production of a bilateral pair of cell clusters as in the embryo. In parallel, we tested the *in vivo* relevance of similar distributions of Cxcl12a/Cxcr4b pathway activation using IR-LEGO ^12^. Interestingly, inducing expression of *cxcl12a* centrally rescues the morphogenesis of the early olfactory epithelium in *cxcl12a* mutant embryos, validating our simulation. Inducing *cxcl12a* expression in an anterior position failed to rescue placode formation, despite our simulations suggesting it should. This discrepancy between *in silico* predictions and *in vivo* observations prompted us to look at the endogenous signaling activity of the Cxcl12a pathway. Our analysis revealed that the chemokine pathway activation dynamics in the anterior compartment are distinct from more posterior regions. Thus, numerically exploration and experimental validation suggest how the distribution of the Cxcl12a signaling drives olfactory placode morphogenesis, and highlighted a spatially controlled dynamic of Cxcr4b activation.

## Results

### Mathematical model to explore the morphogenesis of the olfactory epithelium

We developed a mathematical model (Fig. 1, A and C), to test the contribution of chemotaxis, independently of differentiation, during the formation of the olfactory epithelium on either side of the developing forebrain. We chose a center-based framework, considering each cell as a mobile agent characterized by its position and radius. This allowed access to the dynamics of the system and the flexibility to vary the chemotaxis properties. Center-based models have already been determinant in understanding the emergence of collective organization in complex systems, in particular in the context of chemotactic cell migration ^13–18^.

**Fig 1.**
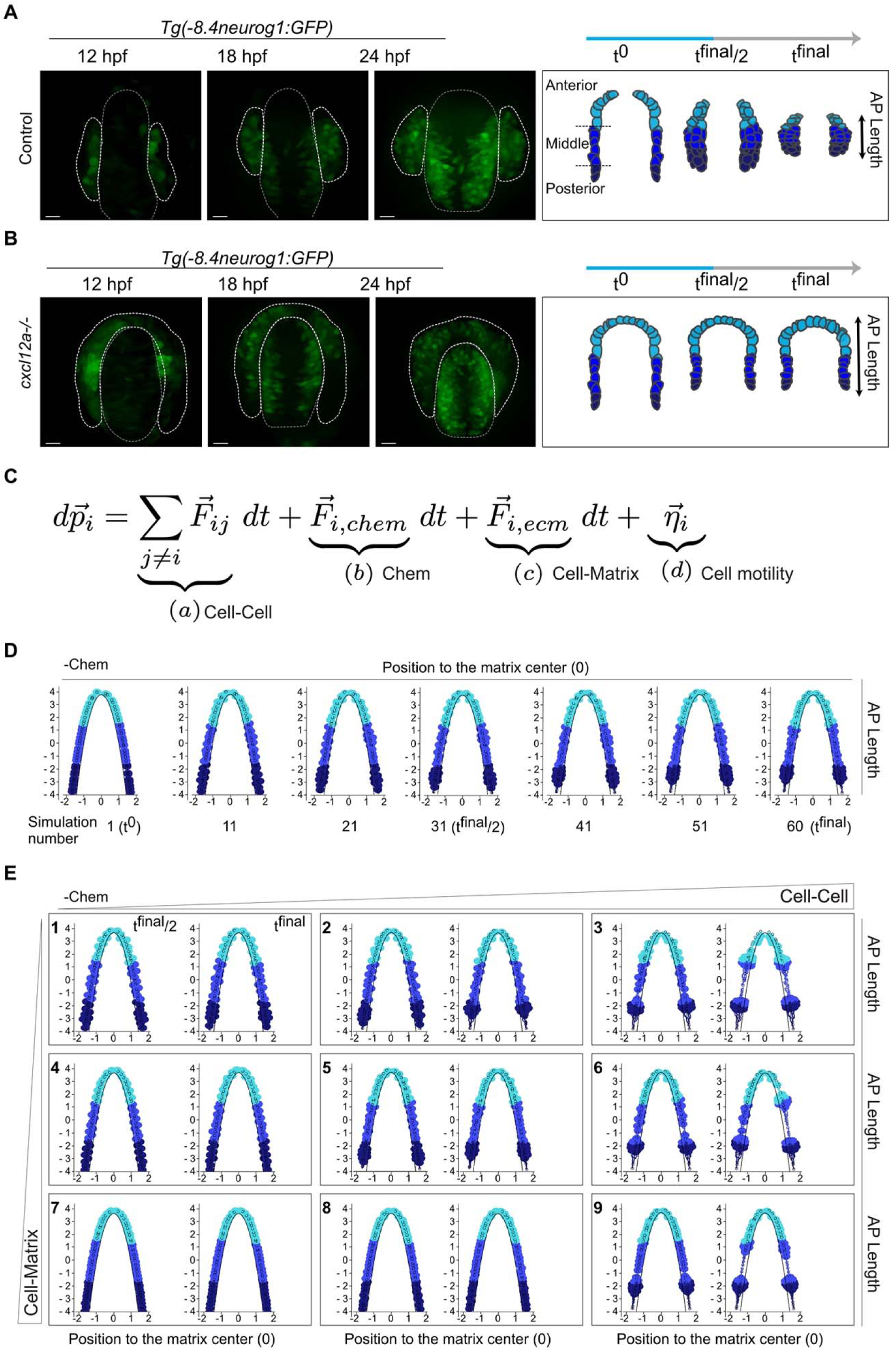
Building a model of olfactory morphogenesis. Visualization of olfactory rosette formation in the olfactory epithelium at 12, 18 and 24 hpf using the *Tg(-8.4neurog1:gfp)^sb1^*line to label pioneer olfactory neurons in control (A), and *cxcl12a^t30516^*mutant (B), conditions. A schema of cell position is shown below with anterior cells shown in light blue, mid-blue and posterior cells in dark blue and the measurement of their size along the anteroposterior (AP) axis is visualized by a black double arrow. Scalebars represent 100 μm. (C), Model equation with the 4 parameters: Cell-Cell interactions (a), Chemotaxis (b), Cell-Matrix interaction (c) and Cell motility (d). (D), Visualization of the positioning of the cells stained in blue scale according to their positions along the AP axis during a simulation in the absence of chemotaxis (-Chem). All 10-time steps are shown. (E), Representation of t^0^ in small box then t^final/2^ and t^final^ of a simulation in the absence of chemotaxis (-Chem) as a function of the intensity of the Cell-Cell and Cell-Matrix forces.

In this model, each cell (i) is represented as a circle of constant radius and characterized by its position p i(t) at each time point t. We considered that cell displacement is defined by 4 main forces: interactions with neighbors (a, Cell-Cell, attraction at short distances and hard-core repulsion), chemo-attraction (b, Chem, toward the source of a chemo-attractant), adhesion to the surface over which it moves (c, Cell-Matrix, short range attraction and hard-core repulsion) and the random motility of an individual cell (d, Cell motility, random motion) (fig. S1A, see “Model details” in Materials & Methods). Considering the system to be at low Reynolds number, we obtained the motion equation for each cell due to the equilibrium of these forces as:

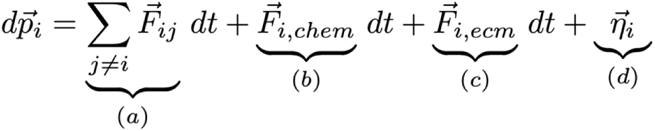

For simplicity, the anterior neural plate was represented as a fixed parabolic, uncrossable barrier for the cells and simulations were performed in 2D with a constant number of cells. The three key stages of olfactory morphogenesis, namely the olfactory territory at 12 hpf, the olfactory placode at 18 hpf and the olfactory epithelium at 24 hpf, are represented by t^0^, t^final/2^ and t^final^ (Fig. 1, A and B). Initially, cells were positioned along the parabolic matrix (20 anterior, 20 middle and 20 posterior). The python code of the simulations is available on github (https://github.com/gletort/Morphoe) and the specific parameters for each figure are also available (https://github.com/gletort/Morphoe/tree/main and https://github.com/JulieBatut/Zilliox-Letort_2024).

To explore the parameters of the model, we first considered the scenario of an absence of chemotaxis (-Chem in the simulations, Fig. 1B). *In vivo*, in the absence of *cxcl12a, Tg(-8.4neurog1:GFP)*-labelled EONs do not converge correctly along the anteroposterior axis, with anterior cells remaining anteriorly positioned ^10^. Figure 1D shows the output of one simulation every 10-time steps in the absence of chemotaxis (*Movie S1*). In this case, anterior cells (light blue) remained in an anterior position, while towards the middle of the simulations, posterior cells (dark blue) migrated slightly anteriorly, reproducing the phenotype described in the absence of *cxcl12a* (Fig. 1D, Frame 31, t^final/2^ and *Movie S1*, fig. S1, B and C-5). We hypothesized that this behavior of posterior cells was generated by the short-range cell-cell attraction with neighboring cells. Supporting this idea, reducing the strength of adhesion between cells, Cell-Cell (Fig. 1E (panels1, 4 and 7), and Movies S2, S5, S7), decreased the displacement of posterior cells. Increasing Cell-Cell augmented cells compaction leading to the presence of several clusters at the end of simulations (Fig. 1E panels 3, 6 and 9, and *Movies S4, S6, S9*). Varying the adhesion of cells to the telencephalon, Cell-Matrix, had a lower impact than varying Cell-Cell (Fig. 1E and *Movies S1-S9*) in the range of values tested. At high values, cells tended to remain arrayed along the parabolic surface, which we will hereafter refer to as matrix, maximizing the overall contact, whereas at lower values, Cell-Cell appears to dominate and favors cluster formation. Thus, in absence of chemotaxis our model predicts that Cell-Cell interactions are the main motor of cell migration.

In simulations, an intermediate strength of Cell-Cell and Cell-Matrix (Fig. 1E, panel 5, *Movie S1*) resulted in a similar distribution of cells to that observed in mutants for *cxcl12a* (^10^, Fig. 1B). To further look at this similarity, we extracted trajectories of the anterior, middle and posterior positioned cells over time (compare fig. S1 B-D with ^10^). The trajectories of the cells in simulations are color coded from t^0^ to t^final/2^ and then, represented in black. The average of the 3 positions is also represented in order to compare the trajectories of cells for each condition (fig. S1C). Another parameter of olfactory morphogenesis at this early stage is the compaction of cells along the AP axis ^7,10,19^, which we measured in the simulations as the evolution of their convergence on the y-axis (which corresponds to the AP axis in the embryo) over time (fig. S1E). All these representations indicate that in the absence of chemotaxis, a balance of Cell-Cell and Cell-Matrix strengths allows the generation of an olfactory epithelium similar to that of a *cxcl12a* mutant *in vivo*.

### Effect of the position of *cxcl12a* expression on the morphogenesis of the olfactory epithelium

Previous studies have shown that *cxcl12a* mRNA is present throughout the anterior neural plate/tube ^8,10^, the future telencephalon. Whether this reflects a requirement for such a pattern of chemokine distribution in the correct positioning of the olfactory epithelium at 24 hpf is unclear. To address this, we simulated the outcome of different distributions of the chemotaxis source in the telencephalon by modulating the position of Chem (Fig. 2, *Movies S10-S14*). In previous work, the repartition of the guidance molecules has been either explicitly modeled taking into account signal production, degradation, cell intake ^20^, considered as a flat distribution of signal ^21^ or following a gradient vanishing away from the source ^22,23^. Here, for simplicity, we considered the chemotaxis signal received by each cell as a spatio-temporal constant force directed toward the chemotaxis source, assuming that the cell is sensing a linear chemotaxis gradient.

**Fig 2.**
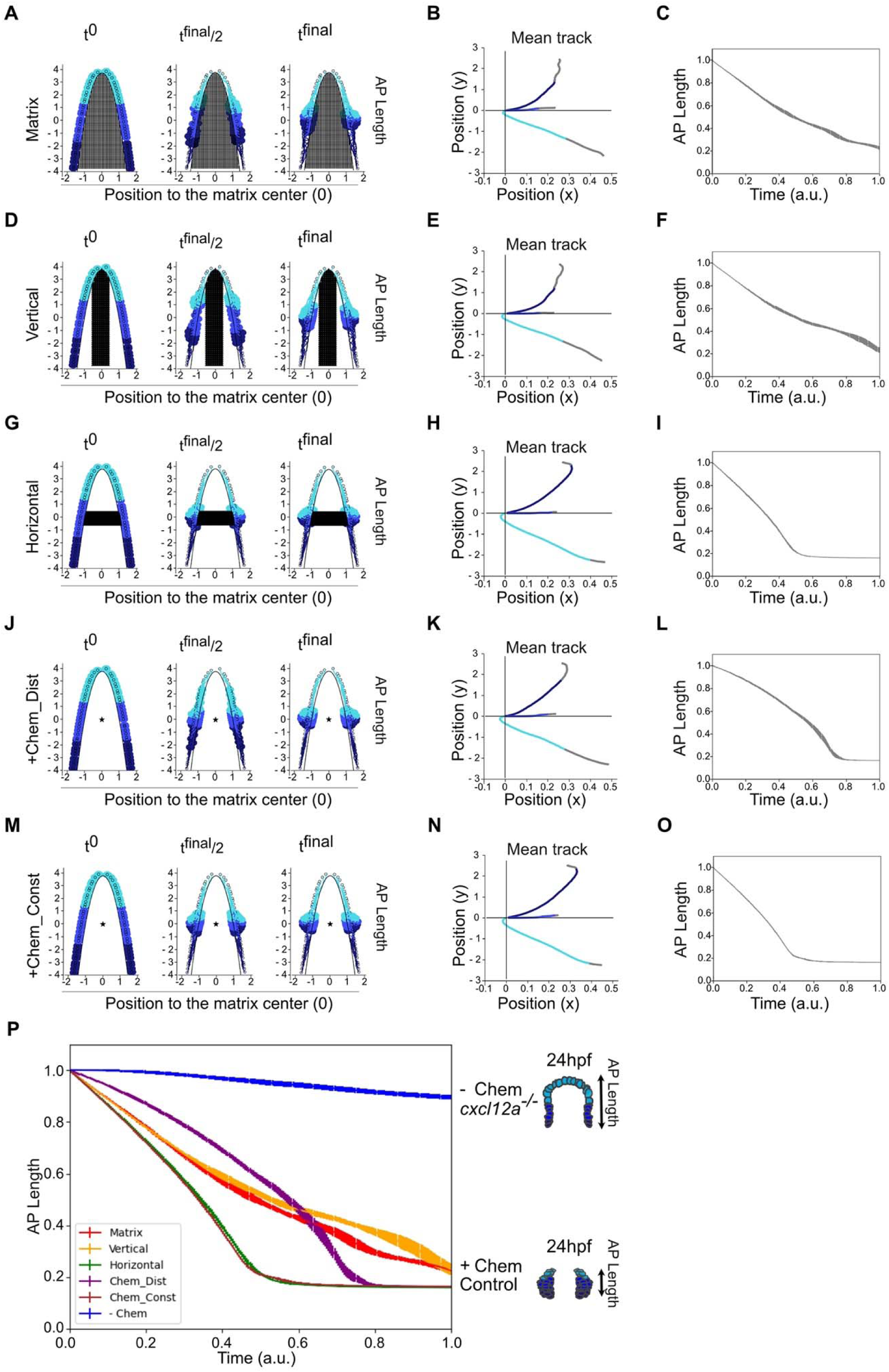
Position of the chemotaxis source controls the compaction of cells along the AP axis to form clusters. Result of a simulation at t^0^, t^final/2^ and t^final^ when the source is (A, Matrix), positioned throughout the matrix, (D, Vertical), vertical, (G, Horizontal), horizontal, (J, +Chem_Dist), a central fixed gradient and (M, +Chem_Const), a central constant force with (B, E, H, K, and N), the representation of the average trajectory of the coded cells as a function of their position on the Antero-Posterior, AP, axis (light blue, anterior; blue, middle and dark blue, posterior). The relative size along the AP axis filled by the cells during the simulations is visualized as a function of the source position (C, F, I, L and 0). (P), Representation of the set of relative sizes along the AP axis occupied by the cells step by step. The curves are colored according to the position of the source (Matrix = Red, Vertical =Yellow, Horizontal = Green, Central source Chem_Dist = Purple and central source, Chem_Const = Brown and without chemotaxis, - Chem = Blue).

We compared the results of cell positioning at t^0^, t^final/2^ and t*^final^* (Fig. 2A, D, J and M), the individual (fig. S2) and average trajectories of 15 simulations of the cells as a function of their position on the AP axis (Fig. 2B, E, K, N) and the convergence of the cells along the AP axis over the course of the different simulations (Fig. 2C, F, I, L, O-P). A chemotaxis source filling the whole matrix (*Movie S10*) or in a vertical/medial position (*Movie S11*) generated similar results but did not correspond to the situation seen in wildtype embryos (Fig. 2A-F and Fig. 2P, red and yellow lines respectively, *Movie S10 and S11*) with, for instance, the size of the cells clusters at t^final^ being too large along the AP axis. We did obtain cell clusters similar to those seen in embryos with simulations based on a Chem source in a horizontal position (Horizontal, Fig. 2G and I, fig. S2C and *Movie S12*). The results are similar with a centrally located point source of Chem (compared Horizontal and Fig. 2G and I and 2J-O as well as the green and brown plots in Fig. 2P, fig. S2C and E and *Movie S14*). Finally, we also tested two scenarios for the evolution of the intensity of the chemotaxis force with a point source: (1) a fixed gradient of intensity (increasing force as cells get closer to the source, +chem_dist (Fig. 2J-L and 2P, *Movie S13*), or (2) a constant force (independent of the distance to the source, +Chem_const, Fig. 2M-P, *Movie S14*). A central source with a chemotaxis intensity gradient (+Chem_Dist) showed correct cluster formation (Fig. 2J, K, *Movie S13*), but not a kinetic evolution of the AP axis length comparable to the control situation in the embryo (^9^ from which fig. S2F is derived).

In conclusion, simulations with a horizontal source of Chem or a point source and a constant force were sufficient to recapitulate the main outcome of olfactory clusters formation.

### Restoring localized source of Cxcl12a rescues the formation of olfactory clusters

According to our simulations, a point source of Chem in the middle of the future telencephalon drives correct morphogenesis of the olfactory epithelium. We tested this *in vivo* by inducing Cxcl12a expression using IR-LEGO (infrared laser-evoked gene operator) ^12,24^ in *Tg(hsp:mcherry-cxcl12a)* transgenic embryos mutant for *cxcl12a* ^25^.

Wild-type embryos formed cell clusters that were visualized by expression of a membrane-bound GFP in a *Tg(cldnb:lyn-GFP)* background, whereas in *cxcl12a* mutant clusters adopted an elongated shape along the AP axis (Fig. 3A). Measurement of the AP axis of each epithelium confirmed these results, with an average of 622 μm ± 25.687 against 801.7 μm ± 19.418 per epithelium, n=7 epithelia from 4 wild-type embryos and 10 epithelia from 5 mutant embryos (Fig. 3C, compared control in blue and mutant in red). IR-induced expression of Cxcl12a in the center of the anterior neural plate at 12 hpf (Fig. 3B, C, *Tg(hsp:mcherry-cxcl12a))* did not affect the AP length of clusters at 24 hpf in the wild-type conditions (642.429 μm ± 7.816 n=7 epithelia, Fig. 3B, C, Blue); expression was visualized by the expression of mCherry (fig. S3). In *cxcl12a* mutant embryos, induced expression of Cxcl12a restored a cluster size comparable to the control situation (688.400 μm ± 20.973, n=10 epithelia, Fig. 3 B, C, Red). Altogether, these data strengthened the predictive capacity of our mathematical model. On the other hand, they open the question of the pertinence of the broad transcription of *cxcl12a* in the presumptive telencephalon, as this configuration of Chem does not produce wild-type convergence of olfactory cell clusters in our simulations.

**Fig. 3.**
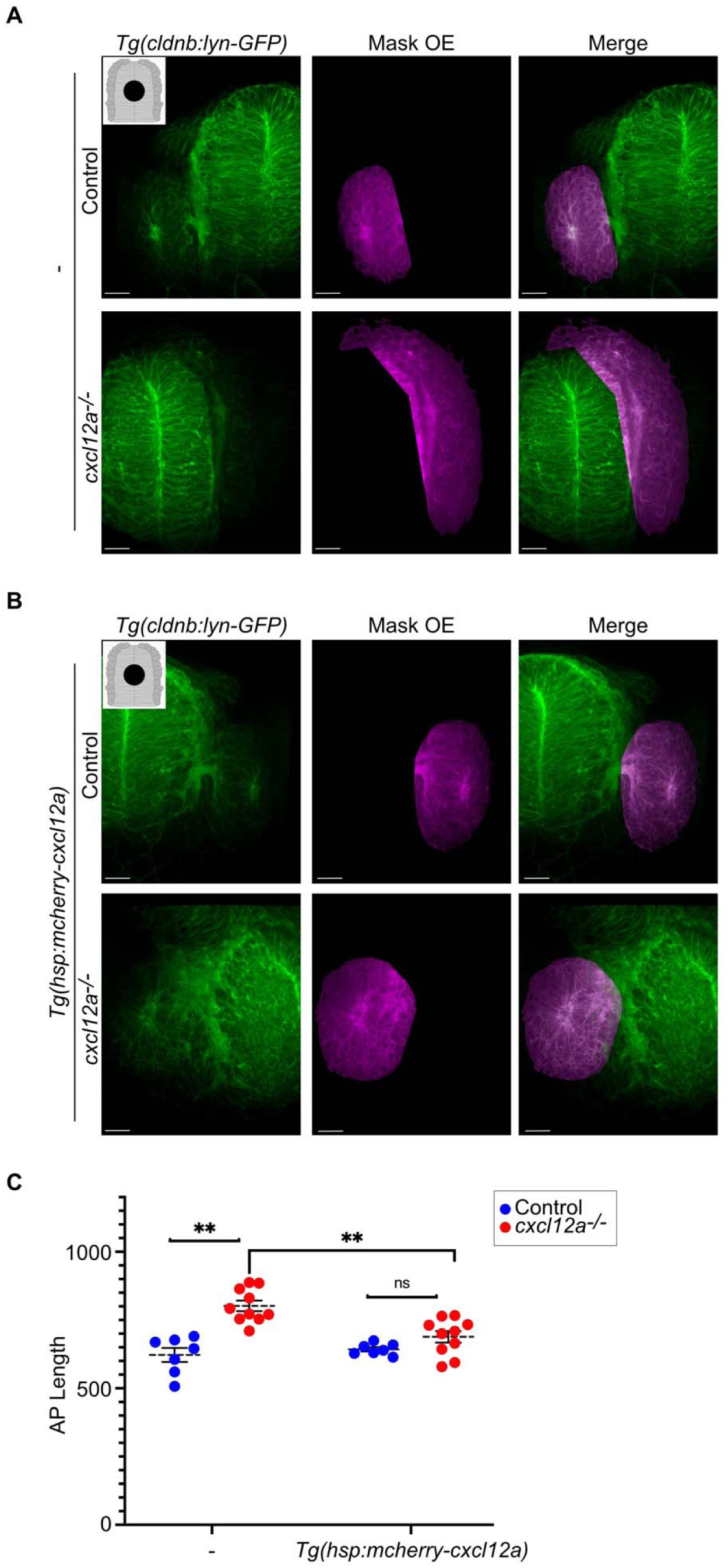
Expression in the middle of the forebrain of *cxcl12a* can rescue olfactory rosette formation in *cxcl12a^t30516^* mutant. (A), Olfactory epithelium (OE) formation visualized at 24 hpf using the *Tg(cldnb:lyn-GFP)* transgenic line in control and *cxcl12a* mutant (*cxcl12a^-/-^)* embryos under control (A, -), or [B, *Tg(hsp:mcherry-cxcl12a*)], conditions in the presence of central *cxcl12a* expression. Rosettes are visualized using the Imaris surface tool and represented by a Mask OE. A merge of the *Tg(cldnb:lyn-GFP)* line and the mask is shown. The embryo is shown with anterior up. Scalebars represent 100 μm. (C), Measurement of the length of the AP axis occupied by cells at 24 hpf from the area occupied by *Tg(cldnb:lyn-GFP)* in control (Control, blue) or *cxcl12a* mutant (*cxcl12a^-/-^*, red) conditions. A significant difference is observed between control and mutant embryos in the control condition (-, without exogenous *cxcl12a* expression), **p=0.0016 (mean epithelial size is 622 μm ± 25.687 versus 801.7 μm ± 19.418, n = 7 and 10, respectively). Conversely, when *cxcl12a* is expressed centrally [*Tg(hsp:mcherry-cxcl12a)*] there is no difference between control and mutant, ^ns^ p= 0,1872 (mean epithelial size is 642.429 μm ± 7.816 versus 688.400 μm ± 20.973 epithelium, n = 7 and 10, respectively). However, there is a difference between the size along the AP axis of the epithelia of mutant embryos in the absence and presence of exogenous cxcl12a expression, **p=0.0070 (801.7 μm ± 19.418 versus 688.400 μm ± 20.973 epithelia, n = 10). Shown are mean ± s.e.m. P-values are calculated using a paired t-test, ns; not significant.

### Clustering of cells along the AP axis is sensitive to the position of a point source of Cxcl12a expression

We further challenged our model by testing cluster formation after induction of Chem/Cxcl12a expression at different positions both in simulations (*in silico*) and embryos (*in vivo*) in -Chem or *cxcl12a* mutant embryos, respectively.

Simulations based on a posterior source led to the formation of two clusters in a posterior position with a normal AP axis length (Fig. 4A-C, fig. S4A and Movie S15). The results of an IR-induced posterior Cxcl12a source in *cxcl12a* mutant embryos look similar to the results of the simulation (Fig. 4D, E and K) with formation of two clusters and an average AP length of 570 μm ± 45.806 (IR-induced Chem), n=4 epithelia compared to 780.333 μm ± 38.394 (non-induced Chem), n=3 epithelia in *cxcl12a* mutants (without Chem); in the wild-type embryos the averages were comparable with or without IR-induce Chem with respectively 536 μm ± 22.275, n=4 epithelia and 497.667 μm ± 48.516, n=3 epithelia. Again, the induction of Cxcl12a expression could be visualized specifically by mCherry expression (fig. S4B, C and fig. S5B, C). Simulations based on anterior Chem resulted in the formation of two clusters in an anterior position or merged clusters across the anterior midline but with clustering kinetics along the AP axis comparable to the control situation (Fig. 4F-H, fig. S5A, Movie S16). In contrast, in *cxcl12a* mutant embryos, anterior expression of Cxcl12a did not rescue the mutant phenotype (Fig. 4I, J and L). After IR-induction clusters displayed an average AP axis length of 795.500 μm ± 55.565, n=4 epithelia while without induction the average AP axis was 718.500 μm ± 34.192, n=4 epithelia; AP axis length in control embryo was 535.333 μm ± 22.615, n=3 epithelia and 460 μm ±63.714 n=4 epithelia, in the absence or presence of induced Cxcl12a expression, respectively (Fig. 4L).

**Fig. 4.**
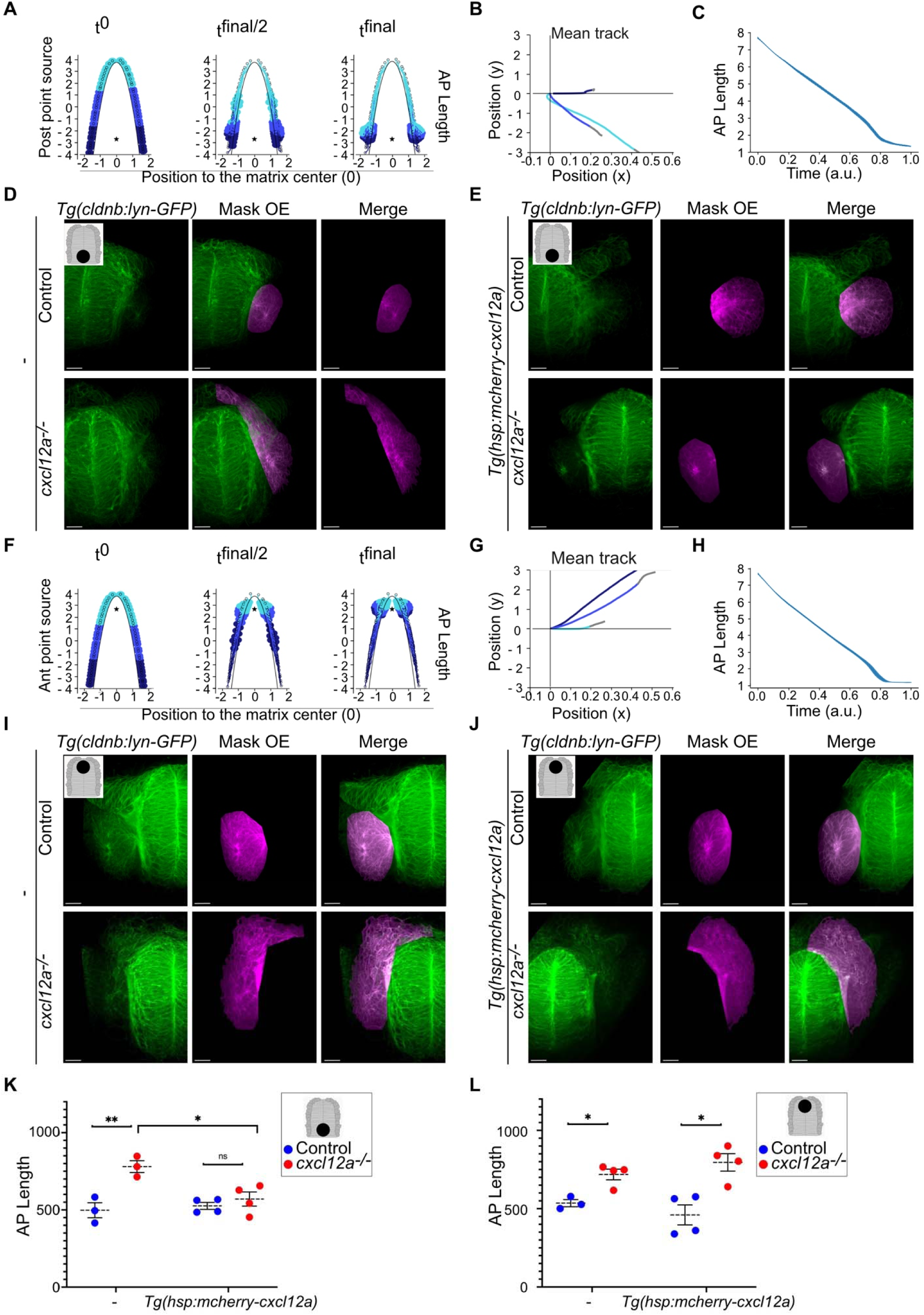
Model and embryo testing of posterior and anterior expression of *cxcl12a* on the morphogenesis of the olfactory epithelium. Simulation of olfactory epithelium formation at t^0^, t^final/2^ and t^final^ in the presence of an (A), posterior or (F), anterior source. (B, G), Average cell trajectory as a function of cell location. Anterior in light blue, middle in blue and posterior in dark blue. (C, H), Time course of the length occupied by the cells along the AP axis. (D), Olfactory epithelium (OE) formation visualized at 24 hpf using the *Tg(cldnb:lyn-GFP)* transgenic line in control and cxcl12a mutant (*cxcl12a^-/-^*) embryos under control (D, -), or [E, *Tg(hsp:mcherry-cxcl12a)*], conditions in the presence of prior *cxcl12a* expression. Rosettes are visualized using the Imaris surface tool and represented by an OE mask. A merge of the *Tg(cldnb:lyn-GFP)* line and the mask is shown. The embryo is shown with the anterior up. Scale bars represent 100 μm. (K), Measurement of the length of the AP axis occupied by cells at 24 hpf from the area occupied by *Tg(cldnb:lyn-GFP)* in control (Control, blue) or *cxcl12a* mutant (*cxcl12a^-/-^*, red) conditions when *cxcl12a* is expressed in the posterior. A significant difference is observed between control and mutant embryos in the control condition (-, without exogenous *cxcl12a* expression), **p=0.0013 (mean epithelial size is 497.667 μm ± 48.516 vs. 780.333 μm ± 38.394, n = 3 and 3, respectively). Conversely, when *cxcl12a* is expressed posteriorly [*Tg(hsp:mcherry-cxcl12a*)], there is no difference between control and mutant, ^ns^p= 0.4849 (mean epithelial size is 526 μm ± 22.275 vs. 570 μm ± 45.806, n = 4 and 4, respectively). However, there is a difference between the size along the AP axis of the epithelia of mutant embryos in the absence and presence of exogenous *cxcl12a* expression, *p=0.0382 (780.333 μm ± 38.394 versus 570 μm ± 45.806 epithelia, n = 3 and 4 respectively). (I), Olfactory epithelium (OE) formation visualized at 24 hpf using the *Tg(cldnb:lyn-GFP)* transgenic line in control and *cxcl12a* mutant (*cxcl12a^-/-^*) embryos under control (I, -), or [J, Tg(*hsp:mcherry-cxcl12a*)], conditions in the presence of anterior *cxcl12a* expression. Rosettes are visualized using the Imaris surface tool and represented by an OE mask. A fusion of the *Tg(cldnb:lyn-GFP)* line and the mask is shown. The embryo is shown with the anterior up. Scale bars represent 100 μm. (L), Measurement of the length of the AP axis occupied by cells at 24 hpf from the area occupied by *Tg(cldnb:lyn-GFP)* under control (Control, blue) or *cxcl12a* mutant (*cxcl12a^-/-^*, red) conditions when *cxcl12a* is expressed in the anterior. A significant difference is observed between control and mutant embryos in the control condition (-, without exogenous *cxcl12a* expression), *p=0.0313 (mean epithelial size is 535.333 μm ± 22.615 vs. 718.500 μm ± 34.192, n = 3 and 4, respectively). Conversely, when *cxcl12a* is expressed anteriorly [*Tg(hsp:mcherry-cxcl12a)*], there is a significant difference between control and mutant, *p= 0.0487 (mean epithelial size is 460.00 μm ± 63.714 vs. 795.500 μm ± 55.565, n = 4 and 4, respectively). Shown are mean ± s.e.m. P-values are calculated using a paired t-test, ns; not significant.

From these results, we conclude that our model faithfully predicts the behavior of inducing central and posterior expression of Cxcl12a in absence of Chem. Conversely, while an anterior point source of Chem generated two anterior clusters in the model, IR-lego Cxcl12a induced anteriorly has no effect in the mutant. This discrepancy between *in silico* and *in vivo* observations challenged our simplistic assumptions, suggesting that activation of the chemokine pathway downstream of the ligand is modulated along the AP axis.

### The anterior part of the olfactory placode shows specific Cxcl12a signaling dynamics

To understand the differences in the predictive power of our model along the AP axis, we analyzed the dynamics of activation of the Cxcl12a pathway in embryos using the previously developed *Tg(cxcr4b-mKate2-IRES-EGFP-CAAX)* transgenic reporter of Cxcl12a signaling ^26^ (fig. S6A, and https://github.com/JulieBatut/Zilliox-Letort_2024/tree/main/Figure5). In this reporter line, activation of the pathway leads to the internalization of Cxcr4b and the appearance of mKate2 puncta in the cytoplasm of cells with EGFP-CAAX+ membranes (blue arrowheads in Fig. 5A and B, and fig. S6). To analyze imaging data, we developed a Java Fiji ^27^ macro (see Materials and Methods) to quantify the number of mKate2 puncta in cells with EGFP-CAAX membranes using Cellpose-based segmentation ^28^. Our results indicate that the dynamics of pathway activation are similar in the posterior and middle compartments, increasing from 12 to 18 hpf before decreased slightly at 24 hpf (fig. S7 A). Indeed, the number of cells displaying puncta in the middle compartment doubled from 12 hpf to 18 hpf, rising from 73 (±23, n=4) at 12 hpf, to 106 (±40, n=2) at 14hpf, 123 (±53, n=2) at 16 hpf, 156 (±43, n=4) at 18 hpf and 117 (±44, n=4) at 24 hpf. Conversely, during the same time window the number of cells in the anterior compartment displaying pathway activation decreased by 2-folds, from 45 (±10, n=4) at 12 hpf, to 37 (±2, n=2) at 14hpf, 25 (±4, n=2) at 16 hpf, 28 (±9, n=4) at 18 hpf and 60 (±28, n=4) at 24 hpf.

**Fig. 5.**
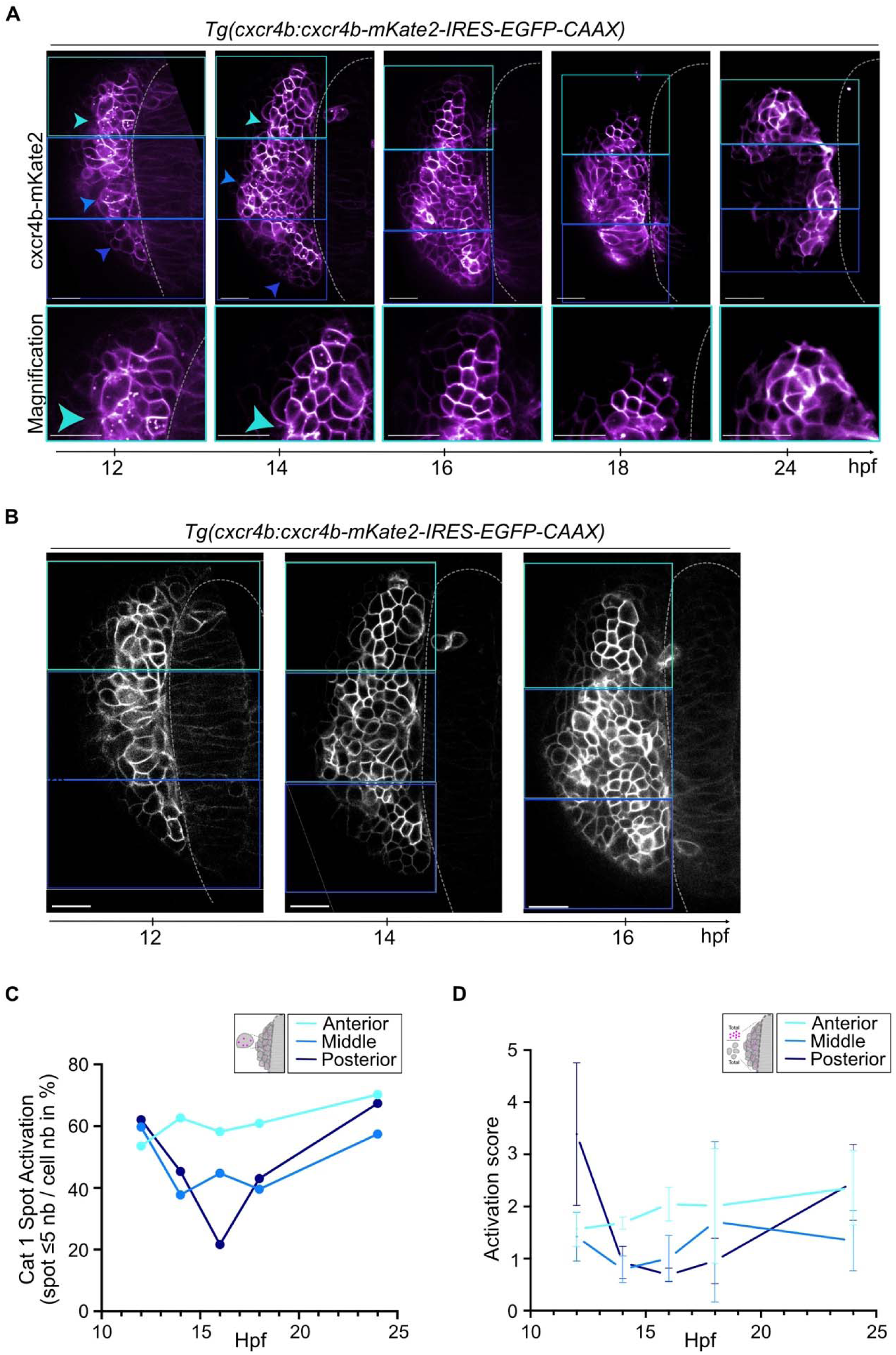
The dynamics of Cxcl12a activation along the AP axis during olfactory epithelium formation. Left placode olfactory cells expressing (A), the Cxcr4b receptor, Cxcr4b-mKate2, and (B), their membranes, EGFP-CAAX, (A, box in bottom right of each image) are visualized over time, yellow arrowheads indicate the Cxcr4b+ spots. (A-B), An optical section is shown. The embryo is shown with the anterior up. Scale bars represent 20 μm. Representation of (C), the percentage of the mean number of intracellular Cxcr4b+ category 1 spots (i.e. percentage of the number of cells with between 1 and 5 Cxcr4b+ spots, relative to the total number of EGFP+ cells, see fig. S5A) and (D), the activation score of the Cxcl12a pathway (total number of Cxcr4b+ intracellular spots over the total number of EGFP+ cells) in anterior (light blue), middle (blue) and posterior (dark blue) cells as a function of time. Shown are mean ± s.d.

**Fig. 6.**
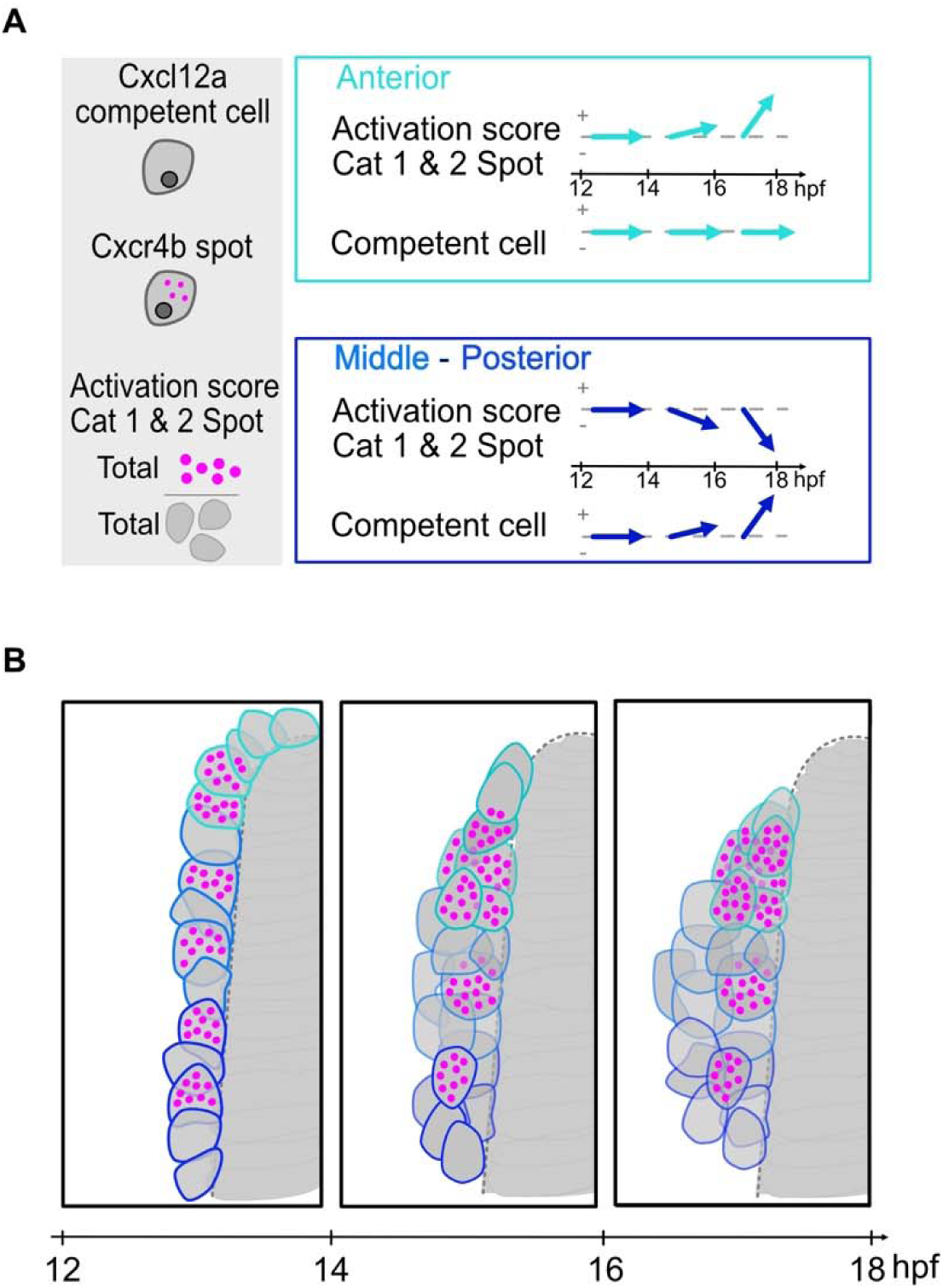
Schematic representation of the activation dynamics of Cxcl12a signaling. (A), Summary of the evolution of the parameters quantified in Fig. 5 and fig. S7 to analyze Cxcl12a signaling. Nuclei are shown in dark gray. Activation of Cxcl12a signaling is visualized by magenta circles in the cytoplasm (light gray). The anterior compartment is shown in light blue while the middle-posterior compartments are shown in dark blue. (B), Schematization of the dynamic evolution from 14 to 18 hpf of activation of Cxcl12a signaling in a left olfactory placode during the compaction process.

Interestingly, we found similar trends when we quantified the number of cells that have Cxcr4b at their membrane but without intracellular spots suggesting that they do not activate the signaling pathway (competent cell, fig. S7 B). The number of competent cells in the middle compartment almost tripled from 30 (±18, n=4) at 12 hpf, to 67 (±36, n=2) at 14hpf, to 68 (±37, n=2) at 16 hpf, to 88 (±25, n=4) at 18 hpf. However, this number remained constant or even decreased slightly in the anterior compartment, from 18 (±8, n=4) at 12 hpf, to 13 (±2, n=2) at 14hpf, to 9 (±4, n=2) at 16 hpf, to 10 (±3, n=4) at 18 hpf (fig. S7B).

We then used two indicators of the presence of Cxcl12a by quantifying the number of intracellular Cxcr4b + points in cells expressing Cxcr4b. Category 1 spots corresponds to the number of cells with 1 to 5 intracellular spots, while Category 2 corresponds to the number of cells with more than 6 intracellular spots relative to the total number of cells. In contrast to the number of competent cells over the same period, the proportion of cells containing between 1 and 5 spots per cell (Cat 1 Spot) remained stable in the anterior cells, whereas it decreased in the other compartments (Fig. 5C). The same trend was observed for cells with a number of Cxcr4b+ spots greater than or equal to 6 (Fig. S7C, Cat 2 Spot). Finally, we determined a Cxcl12a signaling activation score (Fig. 5D) by quantifying the number of total Cxcr4b+ spots over the number of cells expressing Cxcr4b (EGFP-CAAX+). Interestingly, cells from the anterior compartment had an overall higher score than the other compartments, rising from 1.6 (±0.3, n=4) at 12 hpf, to 2 (±0.3, n=2) at 16 hpf, for 1, 4 (±0.5, n=4) at 12 hpf, to 1 (±0.4, n=2) at 16 hpf for cells in the middle compartment and from 3.4 (±1.4, n=4) at 12 hpf, to 0.7 (±0.1, n=2) at 16 hpf for cells in the posterior compartment (Fig. 5D).

Taken together, these results indicate that the anterior part of the olfactory placode has a limited number of cells capable of activating Cxcr4b relative to the middle and posterior compartments. However, activation of this pathway is maintained constant and receptor internalization is higher anteriorly than in other compartments during the early phases of olfactory cell compaction (from 12 to 16 hpf).

## Discussion

Despite increasing knowledge concerning chemotaxis and the control of cell migration, how collectives of cells interpret chemotactic signals during organ formation is still underexplored. Here, we have employed mathematical simulations to help provide insights into this process, focusing on the zebrafish olfactory organ as a model.

We developed a simple mathematical framework to study zebrafish olfactory organ development based on only four parameters: chemoattraction, cell-cell interactions in the collective, cell-matrix interactions and random cell motility. Actual values for the parameters of the model have not been measured. That said, the outcome of a lack of chemoattraction in the system is known as previous studies have shown that embryos mutant for the chemokine Cxcl12a or its receptor Cxcr4b display a characteristic defect in migration of precursor cells that will ultimately form the organ ^8,10^. As such, we were able to set the first parameter of our model to zero and adjust the relative values of those remaining to reproduce the phenotype reported in mutant zebrafish embryos. Our simulations indicated that in the absence of chemotaxis, the force represented by cell-cell interactions is the main driver of cell migration. Consistent with this observation, it has been shown in zebrafish that friction forces at the neuroectoderm-mesendoderm interface are generated by transient E-cadherin-mediated heterotypic contacts between different cell types ^29,30^, reinforcing the idea that cell-cell interactions are central to morphogenesis processes.

With the model appropriately reproducing the *cxcl12a* mutant phenotype, we returned our focus to the chemoattraction parameter. Our simulations showed that cluster formation could be achieved by providing the chemoattractive signal at a variety of positions. Amongst the tested scenarios, however, only a simple gradient of signal from a restricted source induced cluster formation with cell kinetics comparable to those observed in embryos. If this reflects what is indeed happening in the embryo, it implies that the rates of diffusion and degradation of Cxcl12a are relatively constant and that precursors cells of the olfactory organ are competent to respond to the signal independently of their position along the AP axis. Also, and contrary to the broad transcription of *cxcl12a* throughout the anterior neural plate at early stages of development ^8,10^, it suggests that the source of Cxcl12a protein is restricted within the anterior neural plate. This possibility is supported by the *in vivo* rescue of the *cxcl12a* mutant phenotype by IR-LEGO-induced restoration of Cxcl12a expression centrally in the anterior neural plate. An alternative hypothesis to explain our simulations would be that Cxcl12a is produced throughout the telencephalon, in keeping with the transcription of the gene, and that an unknown mechanism in the precursor population would limit activation of the pathway. In the zebrafish posterior lateral line primordium, it has been shown that the interpretation of a uniform Cxcl12a stripe is achieved by the expression of two chemokine receptors, Cxcr4b and Ackr3b, whose localization is restricted within the migrating primordium ^26,31^. To date, however, *ackr3b* expression has not been detected in EONs.

Testing the predictions of the mathematical model *in vivo*, we found that while posterior induction of Cxcl12a restored correct morphogenesis in *cxcl12a* mutants, anterior expression was unable to do so. To understand this discrepancy, we studied endogenous Cxcl12a signaling dynamics in live embryos using a pathway activation reporter ^26^. Our data show that pathway activation displays a similar dynamic in the middle and posterior populations of olfactory placode precursors, with the number of olfactory cells competent to internalize Cxcl12a increasing over a period from 12 to 18 hpf. On the other hand, a general decrease in activation is detected in the anterior compartment during this time window, and a small population of anterior precursors maintains high Cxcr4b internalization. Whether these differences underly those observed between our simulations and IR-LEGO rescue experiments remains to be seen. Unfortunately, it is not possible to determine if anterior IR-LEGO is able to induce a population of cells internalizing high levels of Cxcr4b given the genetic tools available. Nonetheless, it will be interesting to determine if the migratory behavior of this population of anterior precursor cells differs from others in the anterior compartment. As predicted by the interplay between our model and *in vivo* experiments, activation of the pathway appears to be modulated within olfactory placode precursors along the AP axis.

Despite the interesting insights we have gained confronting a theoretical framework and *in vivo* experiments, our study has limitations. First, our mathematical model is extremely simplified. We could have tried to take other parameters such as cell proliferation and apoptosis into account. However, during the period of interest, there are few mitoses ^9^ and the described wave of apoptosis only starts at 24 hpf ^6^. Secondly, in this model, we analyzed the distribution of the source in the telencephalon, whereas hindsight suggests that it will be crucial to understand how olfactory cells perceive the Cxcl12a signal. Thirdly, our model does not take into account the evolution of the relationship between cells. A ‘neighborhood watch’ model has been employed to explain the formation of the primitive streak in the chicken embryo because embryonic epiblast cells evaluate positional information relative to their neighbors ^32^. We based our approach on a region-based view of the dynamics of Cxcl12a signaling given out previous work ^10^ but our model lacks an *in vivo* cellular scale. Knowing how the dynamics of Cxcl12a signaling evolves at the cellular scale would give us a better understanding of how cells integrate this signaling in relationship to each other. Finally, our model treats our 4 parameters as independent whereas there are potential links between them, in particular between Cxcl12a signaling and Cell-Cell or Cell-Matrix interactions. Indeed, in the lateral line, all the cells of the primordium must detect the attractant and adhere to each other to coordinate their movements and display a robust directed migration ^33^. This suggests that a more complex, probably multi-scale model will need to be developed in the future to explore the spatial and temporal heterogeneity of Cxcl12a signaling and cell activation further.

## Materials and Methods

Model (https://github.com/JulieBatut/Zilliox-Letort_2024.git).

To explore numerically the formation of the olfactory placode, we modeled the motion of cells at the individual cell level and used an agent-based approach ^34^. For simplicity, cells were considered identical and represented as circles of constant radius that can move and interact. Similar off-lattice particle/cell-centered models were indeed used to simulate multi-cellular cell systems and collective motion ^35–39^. Each cell *i* was represented by the position of its center *p_i_*. Under the standard assumption of low Reynolds number regime, the equation of motion for each cell center was 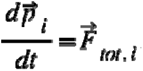

In our model, we considered that the motion of the cells was determined by 4 main factors: the interaction with its neighbors (a), the attraction from the chemotaxis source (b), the interaction with the extra-cellular matrix (telencephalon) (c) and the spontaneous motility of an individual cell (d). Thus, the displacement of one cell *i* was calculated as:

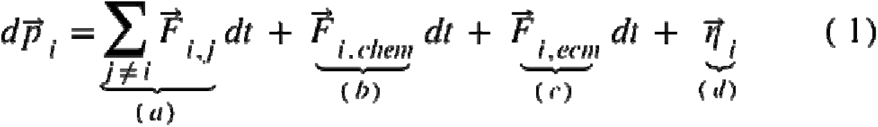

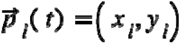 is the position of the center of the cell *i* at time *t*.

### Cell description

For simplification, we described an individual cell as a circle of radius *d_eq_/*2, equal for all cells. However, the cells could interact with each other until a distance *d_lim_* greater than *d_eq_* to take into account the presence of protrusions in the cells (like filopodia).

### Cell-cell interactions (a)

When 2 cells are within a distance smaller than *d_lim_* from each other, their contact will influence their motion. At first, close cells which do not overlap will attract each other by forming junctions upon contact, which permits cells clustering. However, as the cell cannot physically overlap, if the distance between their center decreases below the equilibrium distance *d_eq_* then there will be a hard-core repulsion to account for the limited deformability of the cells.

The force applied on one cell *i* by its neighboring cell *j* is then calculated as:

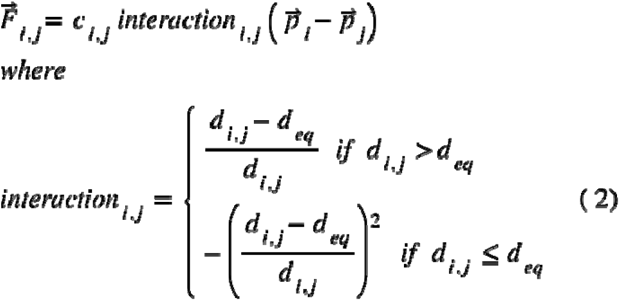

### Chemotaxis (b)

We wanted to test the contribution of a chemotaxis agent present in the middle of the telencephalon. Thus, we added a force attracting the cells toward the point source of coordinates (0,0). We tested two scenarios. In the first one we assume that this attraction depends on the distance of the cell to this point source (4 - dist), and increase as the cell got closer to this point (the cell receives more signal as the cell gets closer to the source). In the second case, the attraction follows a constant gradient (4 - const) and is thus independent from the distance to the source.

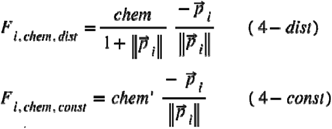

where chem and chem are the strengths of the attraction force in the different cases.

### Cell-matrix interaction (c)

For simplicity, the matrix (telencephalon) was implemented with a parabolic shape, which was sufficient to roughly resemble biological shape. As or the cell-cell interaction, we considered in the model the effect of the interaction between each cell and the matrix. Cells tend to adhere to the matrix which is represented in the model by an attractive force, but cannot cross it, which is calculated as repulsive force between the matrix and the cell. This interaction is thus calculated as:

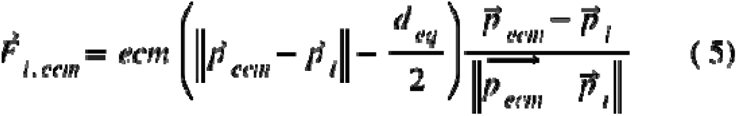

Where ecm is the strength of the attractive/repulsive forces between one cell and the matrix, and 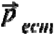 is the closest point of the matrix to the cell (projected point) where the interaction is considered to take place. For numerical reason, we increased the repulsion coefficient if the cell got inside the matrix 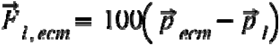 to ensure that the cells stay outside.

### Cell motility (d)

The goal of the model is to test the effect of the different forces on the cell motility, taking into account only essential components. Thus, we simplified the description of the cell motility as a random Brownian motion:

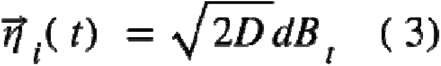

Note that we also tested a model in which we took into account the polarized motion of the cell by keeping a persistence of direction (due to the internal organization of the cell), and that did not impact the general behavior.

All the parameters for each simulation are shown here: https://github.com/JulieBatut/Zilliox-Letort_2024.The source code for the simulations, developed in python are publicly available here: https://github.com/gletort/Morphoe/tree/main

|  |  |
| --- | --- |
| $N$ | 60 |
| $dt$ | 0.0005 |
| $t_{max}$ | 6 |
| $d_{lim}$ | 1 |
| $d_{eq}$ | 0.5 |
| $Chem$ | 2 |
| $D$ | 0.001 |
| $v_0$ | 0.1 |
| $c_{ij}$ | 0.7 |
| $ecm$ | 4 |

#### Fish Husbandry and lines

Ethics Statement and Embryos: All zebrafish and embryos were handled according to relevant national and international guidelines. French veterinary services and the local ethical committee approved the protocols used in this study, with approval ID: E31555011 and APAPHIS #34368-2021121409357964-v6.

Fish were maintained at the CBI zebrafish facility in accordance with the rules and protocols in place. The *cxcl12a^t30516^*mutant line has previously been described ^40^, as have the *Tg(-8.4neurog1:gfp)^sb1^* ^41^, *Tg(-8.0cldnb:lynGFP)^zf106^* ^42^, *TgBAC(cxcr4b:cxcr4b-mKate2-IRES-EGFP-CAAX,cryaa:DsRed)^sk79^* ^26^ in this paper referred as *Tg(cxcr4b-mKate2-IRES-EGFP-CAAX)* and *Tg(hsp70l:mCherry-cxcl12a)^zf3312^*^25^ here referred as *Tg(hsp:mcherry-cxcl12a)*. Embryos were obtained through natural crosses and staged according to ^43^.

#### IR-LEGO and time-lapse spinning disk datasets

Embryos carrying both *Tg(-8.0cldnb:lynGFP)^zf106^* and *Tg(hsp:mcherry-cxcl12a)* transgenes and control or mutant for *cxcl12a^t30516^*were selected. Embryos were then grown to 12 hpf at which point they were dechorionated and embedded for imaging in 0.7% low-melting point agarose in embryos medium. For Infra-Red (IR) LEGO, a 40x HC PL IRAPO/1.10 W CORR lens (Leica, 15506352) was used to perform 90% IR 1470 nm laser illumination for 1min over a 22 μm radius in the middle, anterior or posterior of the telencephalon depending on the experiment. This illumination was performed using the in-house IR-LEGO configuration ^12^ and a time-lapse series of confocal stacks (0,7 μm slice/100 μm deep) was generated of the anterior neural plate and flanking non-neural ectoderm on an inverted Leica DMi8 Spinning microscope using a 40x HC PL APO/1.3 oil objective (Leica, 11506329). Acquisitions each 1 hour were stopped at 24 hpf (t^final^), when the olfactory rosette was clearly visible. The shape of the olfactory rosette was them defined using Imaris 8.3 analysis software (Bitplane, Switzerland). IR-LEGO and Imaging data have been collected by Rapp’Opto software and Metamorph software (version: Metamorph 7.8.13.0) respectively.

#### Cxcl12a dynamic activation live imaging: data collection and processing

Embryos carrying *Tg(cxcr4b:cxcr4b-mKate2-IRES-EGFP-CAAX)* transgene were selected. Embryos were then grown to 12 hpf at which point they were dechorionated and embedded for imaging in 0.7% low-melting point agarose in embryos medium. A time-lapse series of confocal stacks (0,3 μm slice/120 μm deep) was generated of the anterior neural plate and flanking non-neural ectoderm on an inverted Leica DMi8 Spinning microscope using a 40x HC PL APO/1.3 oil objective (Leica, 11506329). Acquisitions each 1 hour were stopped at 24 hpf (t^final^), when the olfactory rosette was clearly visible. A Fiji macro has been developed to analyze Cxcl12a activation in olfactory cells. Briefly, after selection and (anisotropic) filtering of the red (561, Cxcr4b-mKate2) and green (EGFP-CAAX) images, the pools of cells with a green membrane were segmented and extracted using cellpose 2.0 (https://www.cellpose.org/). Finally, a macro to quantify the number of red clusters and their intensities (reflecting Cxcl12a activation by Cxcr4b) was generated. All the data for this quantification are available on github upon request (https://github.com/JulieBatut/Zilliox-Letort_2024.git).

### Statistical analysis

All statistical comparisons are indicated in figure legends including one sample and paired t-test performed using Prism (GraphPad.) The scatter dot plots were generated with Prism. Data are mean ± s.e.m. Two-tailed t-test ^ns^p<0.1234, *p<0.0332, **p<0.0021, ***p<0.0002. For image analysis, data shown are mean ± s.d.

## Acknowledgments

This work was supported by the Centre National de la Recherche Scientifique (CNRS) as part of its *Biomimetisme* interdisciplinary programme over the 2019-2020 period; Université de Toulouse III (UPS); the Ministère de la Recherche and the *Agence Nationale de la Recherche* (ANR), with the ZOORRO grant, ANR-23-CE45-0021. We thank Holger Knaut, Darren Gilmour, Kristen Kwan and Chi-Bin Chien for providing plasmids, Stéphanie Bosch, Brice Roncin, Vanessa Dougados, Thomas Mangeat and the Toulouse RIO Imaging platform, and zootechnicians for taking care of the fish. We also thank EMBO Workshop MMM2017 and members of the Blader and Batut labs for advice and comments on the manuscript.

## Author contributions

Conceptualization: JB

Methodology: MZ, GL, PD, AP, DS, JB

Investigation: MZ, GL, PB, VRM, DS, JB

Visualization: CR, MZ, GL, DS, MZ, JB

Supervision: DS, PB, JB

Writing—original draft: JB, MZ, GL

Writing—review & editing: MZ, GL, FG, PB, DS, JB

## Supplementary Materials

### Supplementary Text

All the movie shows the entire simulation process from t^0^ to t^final^ (1 to 60 frames). https://github.com/JulieBatut/Zilliox-Letort_2024.git (request access if necessary).

---

**Fig. S1.**
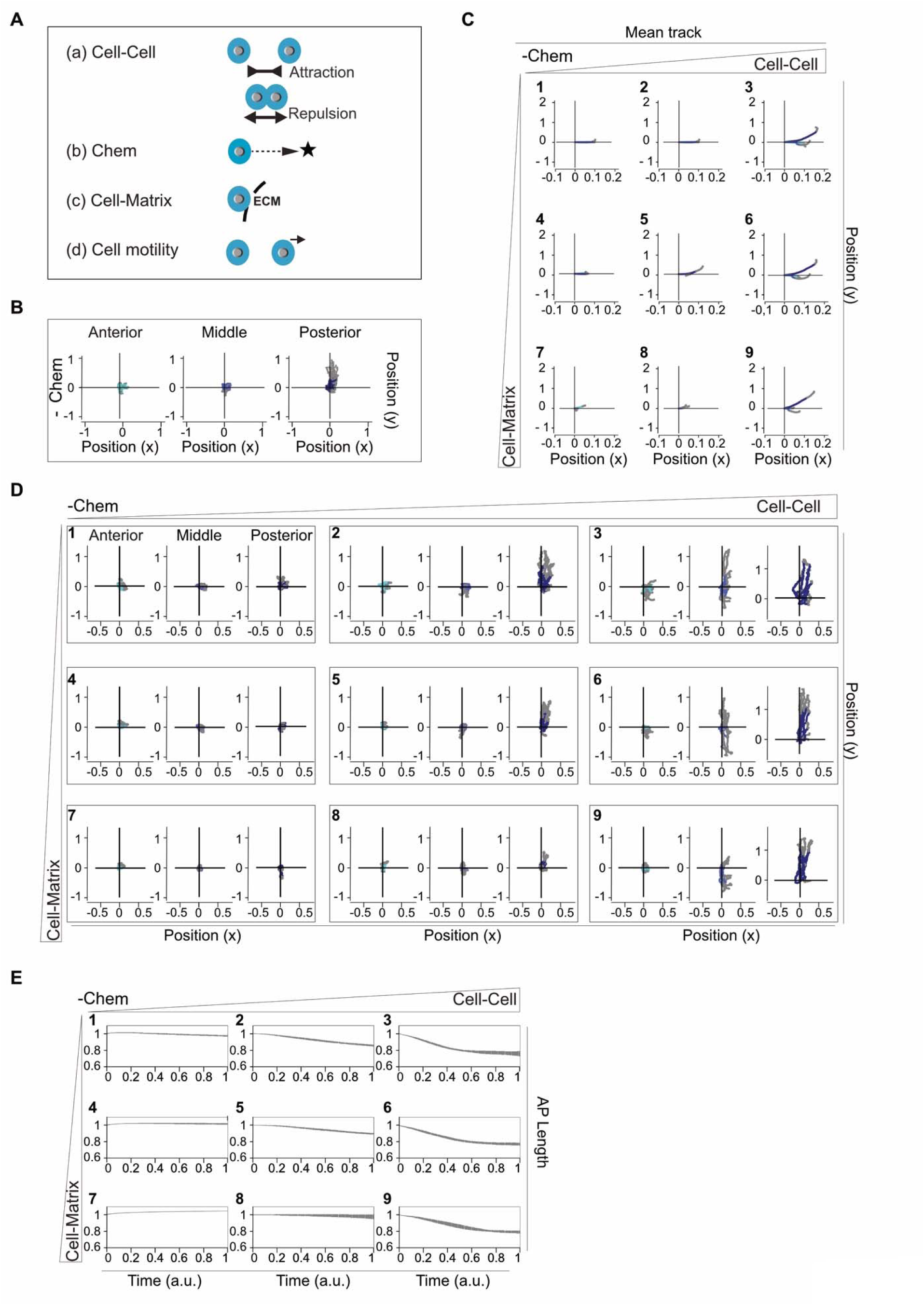
Model parameters to reproduce the morphogenesis of the olfactory epithelium in the absence of chemotaxis *i.e.* to phenocopy the one of the *cxcl12a* mutant. (A), Representation of the 4 parameters of the model and example on a cell represented in blue with a core in grey. The position of the chemotaxis source is visualized by a black star. Chem, chemotaxis; ecm; extra cellular matrix. (B), Individual cell trajectories according to their position in the anterior posterior (AP) axis in simulation in the absence of chemotaxis (-Chem). Representation of the average trajectory of 15 simulations (C), or each trajectory of the anterior, middle and posterior cells (D), as a function of the intensity of the Cell-Cell and Cell-Matrix strength. (B-D). The paths are represented in color (shade of blue according to the position along the AP axis) from t^0^ to t^final/2^ then in gray until t^final^. (E), Representation of the variation in cell size along the AP axis during simulations in the absence of chemotaxis and as a function of the intensity of the Cell-Cell and Cell-Matrix force.

**Fig. S2.**
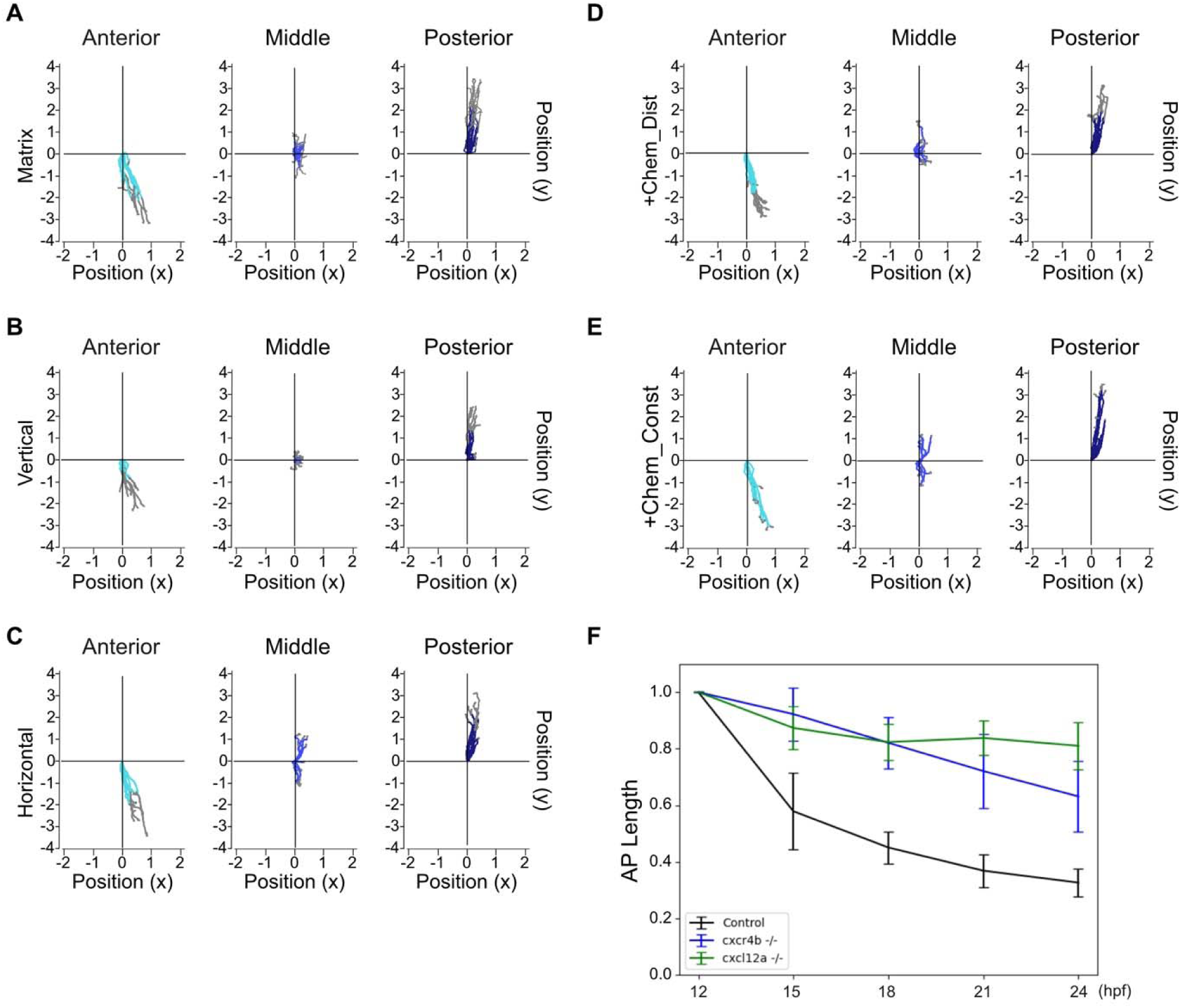
Average trajectories of the cells during the different simulations of the source positions and evolution of the size along the AP axis occupied by the cells during the formation of the olfactory rosettes in the embryo (modified from ^1^). Representation of the average trajectory of the anterior, middle and posterior cells as a function of the positioning of the source, *i.e.* (A), in the whole matrix, (B), vertically, (C), horizontally, (D, +Chem_Dist), with a gradient fixed at the centre or (E, +Chem_Const), with a central constant force. The trajectories are represented in colour (shade of blue depending on the position along the AP axis) from t^0^ to t^final/2^ and then in grey until t^final^. (F), Representation of the relative evolution of the size along the AP axis occupied by the cells every 3 h during the olfactory rosette formation process from 12 hpf to 24 hpf.

**Fig. S3.**
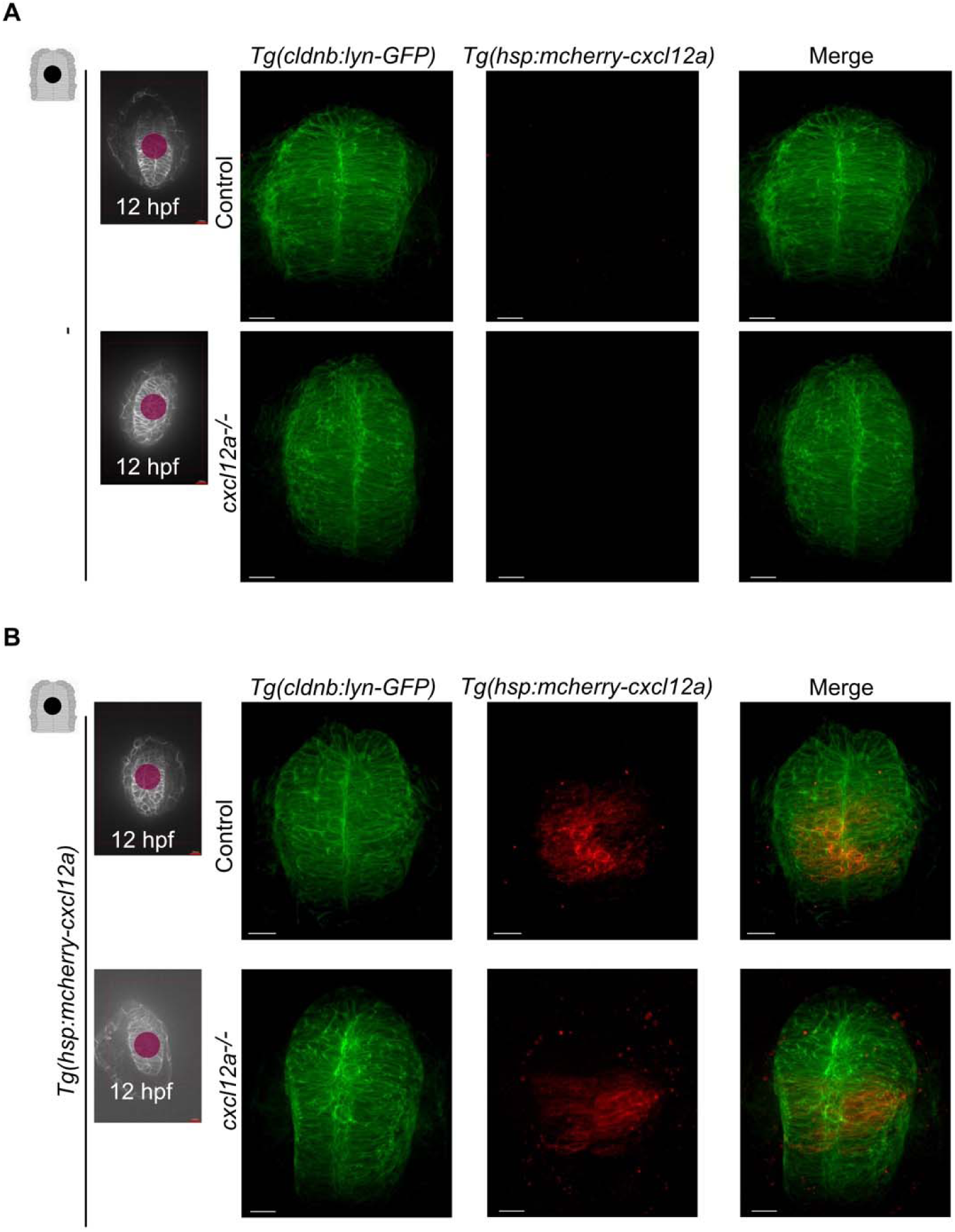
Validation of *cxcl12a* expression in the middle of the telencephalon using the IR-LEGO technique. Visualization of the telencephalon at 16 hpf using the *Tg(cldnb:lyn-GFP)* transgenic line in control and *cxcl12a* mutant *(cxcl12a^-/-^)* embryos after infrared (IR) irradiation in the mid-telencephalon at 12 hpf in the absence (A, -), or in the presence of the [B*, Tg(hsp:mcherry-cxcl12a)* transgene]. (A, B), The IR laser target at 12 hpf is shown in the left panels and a combined red and green channel is also shown (Merge). The embryo is shown with the anterior up. Scale bars represent 100 μm. Following irradiation, (A, -), in the absence of the transgene, there is no expression of *cxcl12a* whereas in the presence of the transgene [B, *Tg(hsp:mcherry-cxcl12a)*], the expression of *cxcl12a* is validated in red in the mid-telencephalon.

**Fig. S4.**
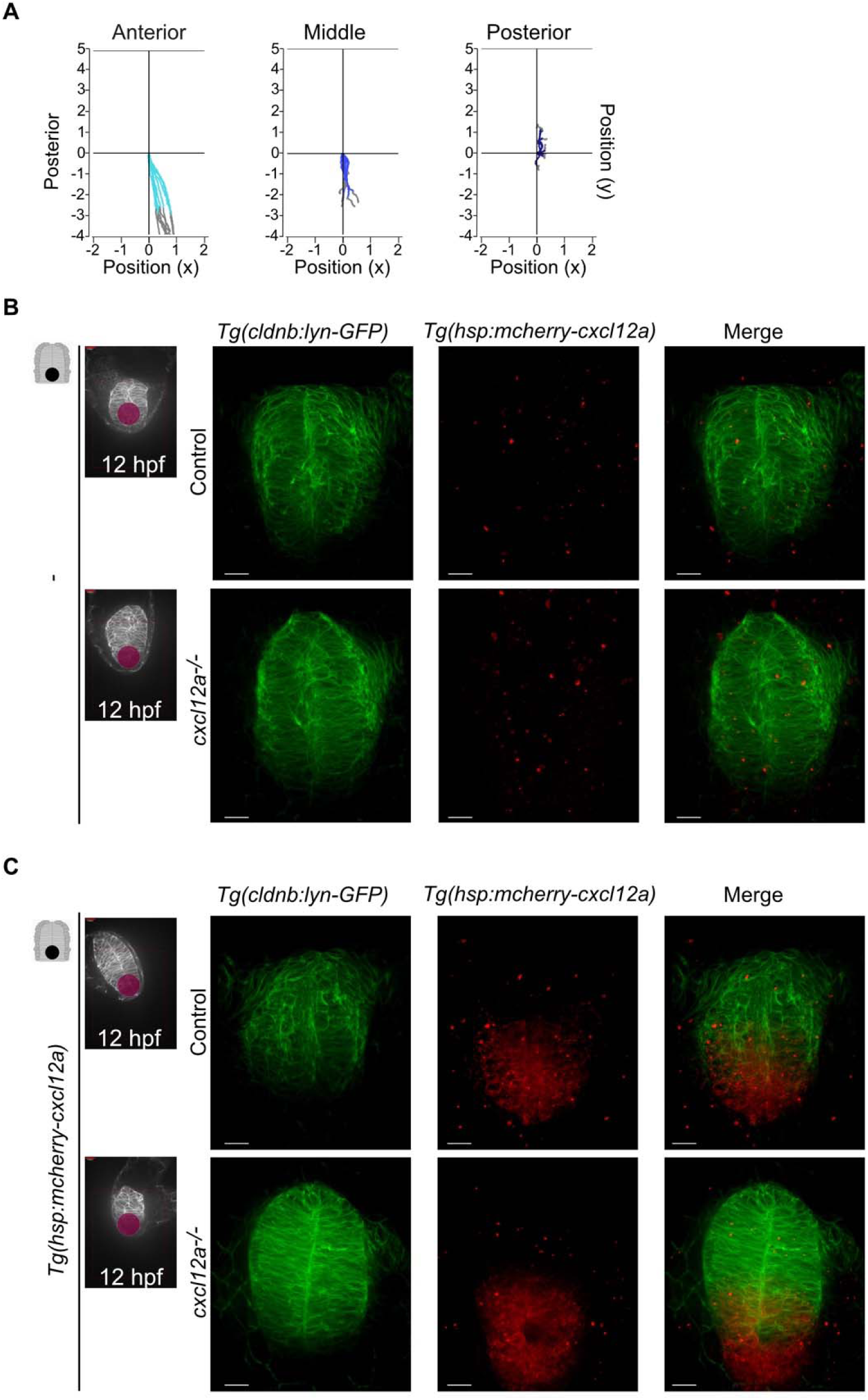
Simulation and validation of *cxcl12a* expression in the posterior telencephalon using the IR-LEGO technique. (A), Representation of the average trajectory of coded cells as a function of their position on the Antero-Posterior axis, (light blue, Anterior; blue, Middle and dark blue, Posterior). Visualization of the telencephalon at 16 hpf using the *Tg(cldnb:lyn-GFP)* transgenic line in control and *cxcl12a* mutant (*cxcl12a^-/-^)* embryos after infrared (IR) irradiation in the posterior part of the telencephalon at 12 hpf in the absence (B, -), or presence (C, *Tg(hsp:mcherry-cxcl12a*), of the transgene. (B, C), The IR laser target at 12 hpf is shown in the left panels and a combined red and green channel is also shown (Merge). The embryo is shown with the anterior up. Scale bars represent 100 μm. After irradiation, (B, -), in the absence of the transgene, no expression of *cxcl12a* is present whereas in the presence of the transgene [C, *Tg(hsp:mcherry-cxcl12a)*], *cxcl12a* expression is validated in red in the posterior telencephalon.

**Fig. S5.**
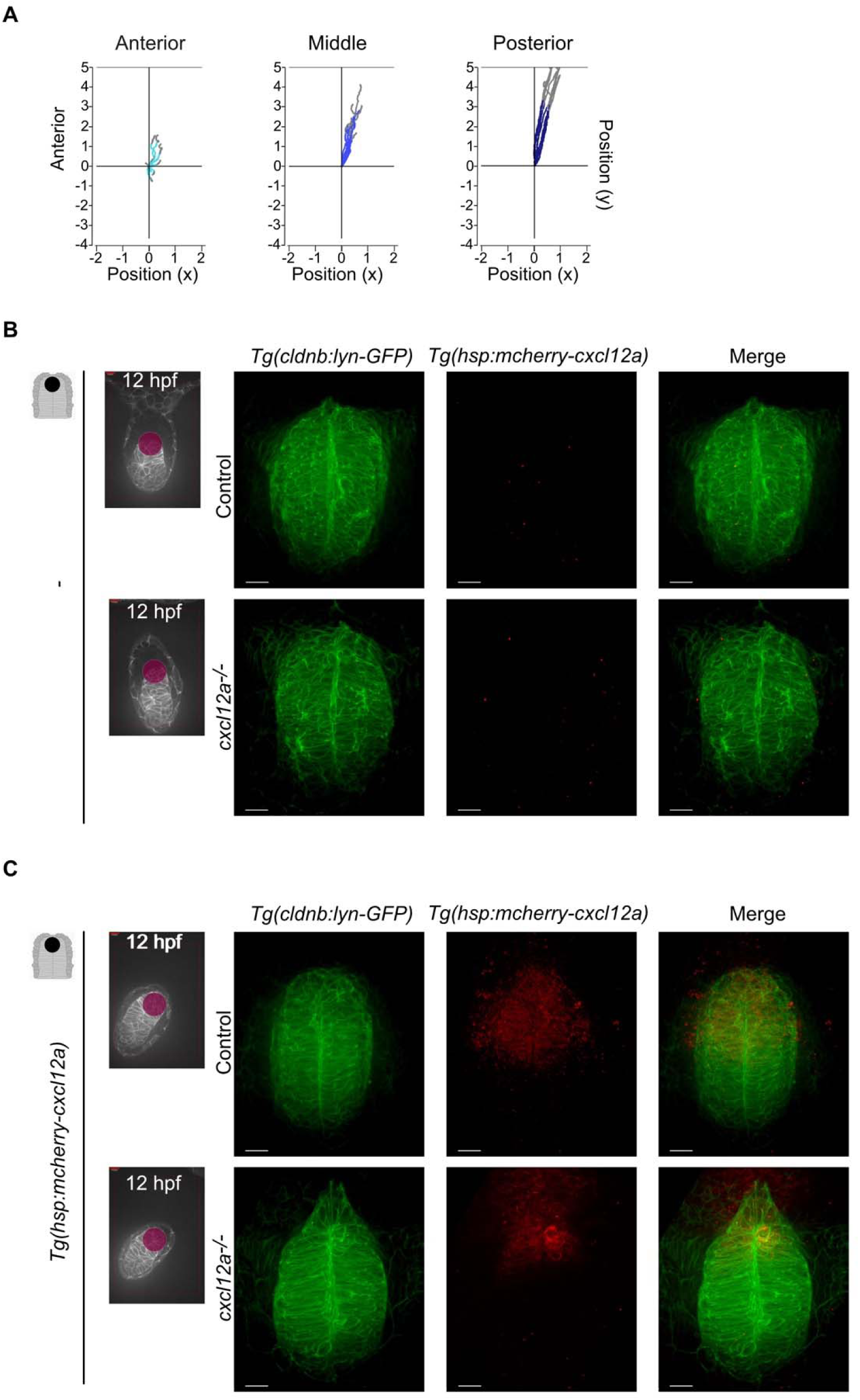
Simulation and validation of *cxcl12a* expression in the anterior telencephalon using the IR-LEGO technique. (A), Representation of the average trajectory of coded cells as a function of their position on the Antero-Posterior axis, (light blue, Anterior; blue, Middle and dark blue, Posterior). Visualization of the telencephalon at 16 hpf using the *Tg(cldnb:lyn-GFP)* transgenic line in control and *cxcl12a* mutant (*cxcl12a^-/-^)* embryos after infrared (IR) irradiation in the anterior part of the telencephalon at 12 hpf in the absence (B, -), or presence [C, *Tg(hsp:mcherry-cxcl12a*)] of the transgene. (B, C) The IR laser target at 12 hpf is shown in the left panels and a combined red and green channel is also shown (Merge). The embryo is shown with the anterior up. Scale bars represent 100 μm. After irradiation, (B, -), in the absence of the transgene, no expression of *cxcl12a* is present whereas in the presence of the transgene [C, *Tg(hsp:mcherry-cxcl12a)*], *cxcl12a* expression is validated in red in the anterior telencephalon.

**Fig. S6.**
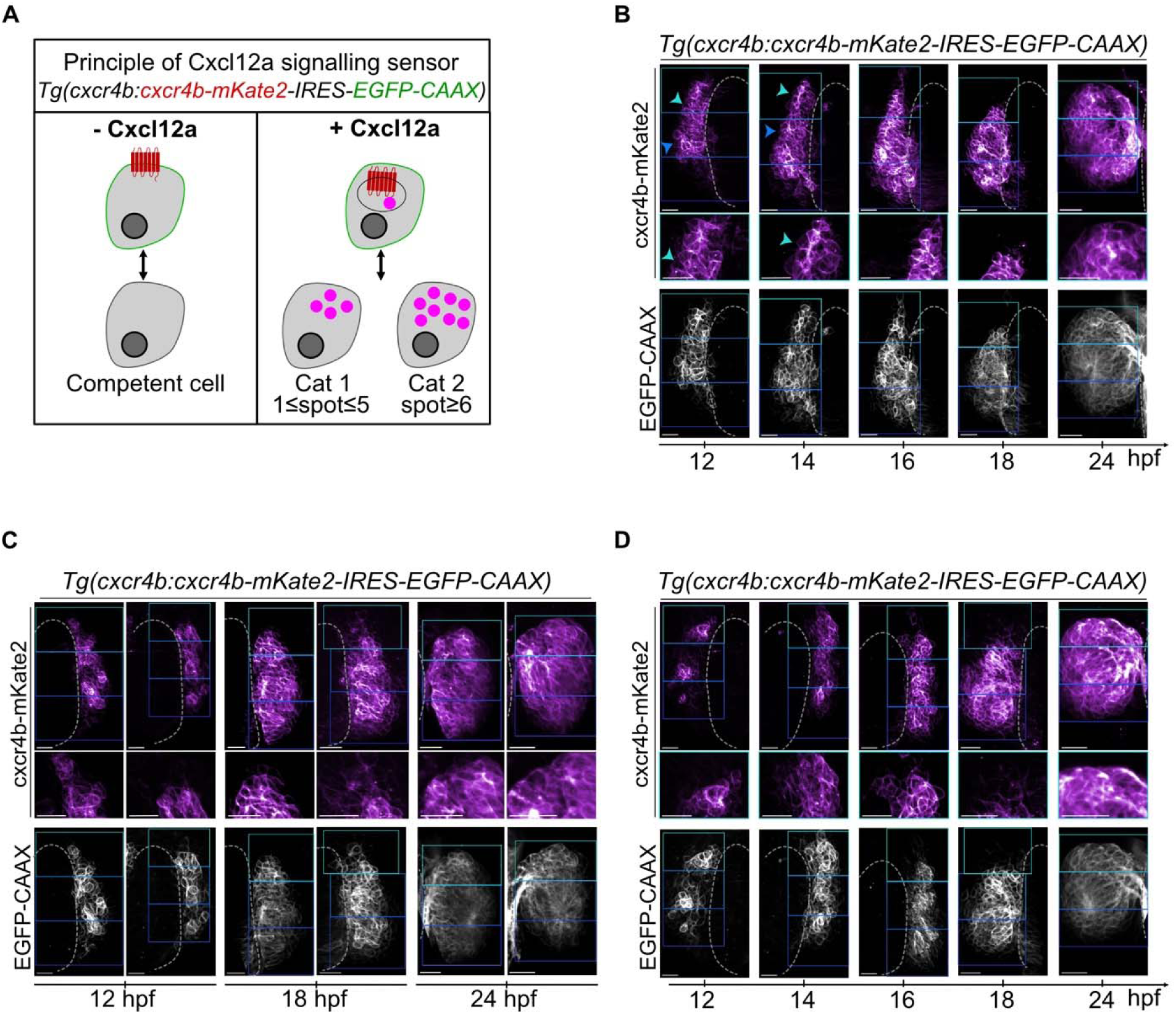
Cxcl12a activation along the AP axis during olfactory epithelium formation. (A), Principle of Cxcl12a signaling sensor in absence (-Cxcl12a) or in presence (+ Cxcl12a) of Ligand. Cxcr4b-mKate2 receptor is shown in magenta. (B), Left or (C), right or (D), left and right placode olfactory cells expressing the Cxcr4b receptor, cxcr4b-mkate2, and their membranes, EGFP-CAAX, visualized over time. The anterior, middle and posterior regions of interest are visualized by colored rectangles with progressively darker levels of blue towards the posterior. Blue arrowheads (with increasing levels of blue towards the posterior) indicate the Cxcr4b+ spot. (B-C, D), An optical section is shown. The embryo is shown with the anterior up. Scale bars represent 20 μm.

**Fig. S7.**
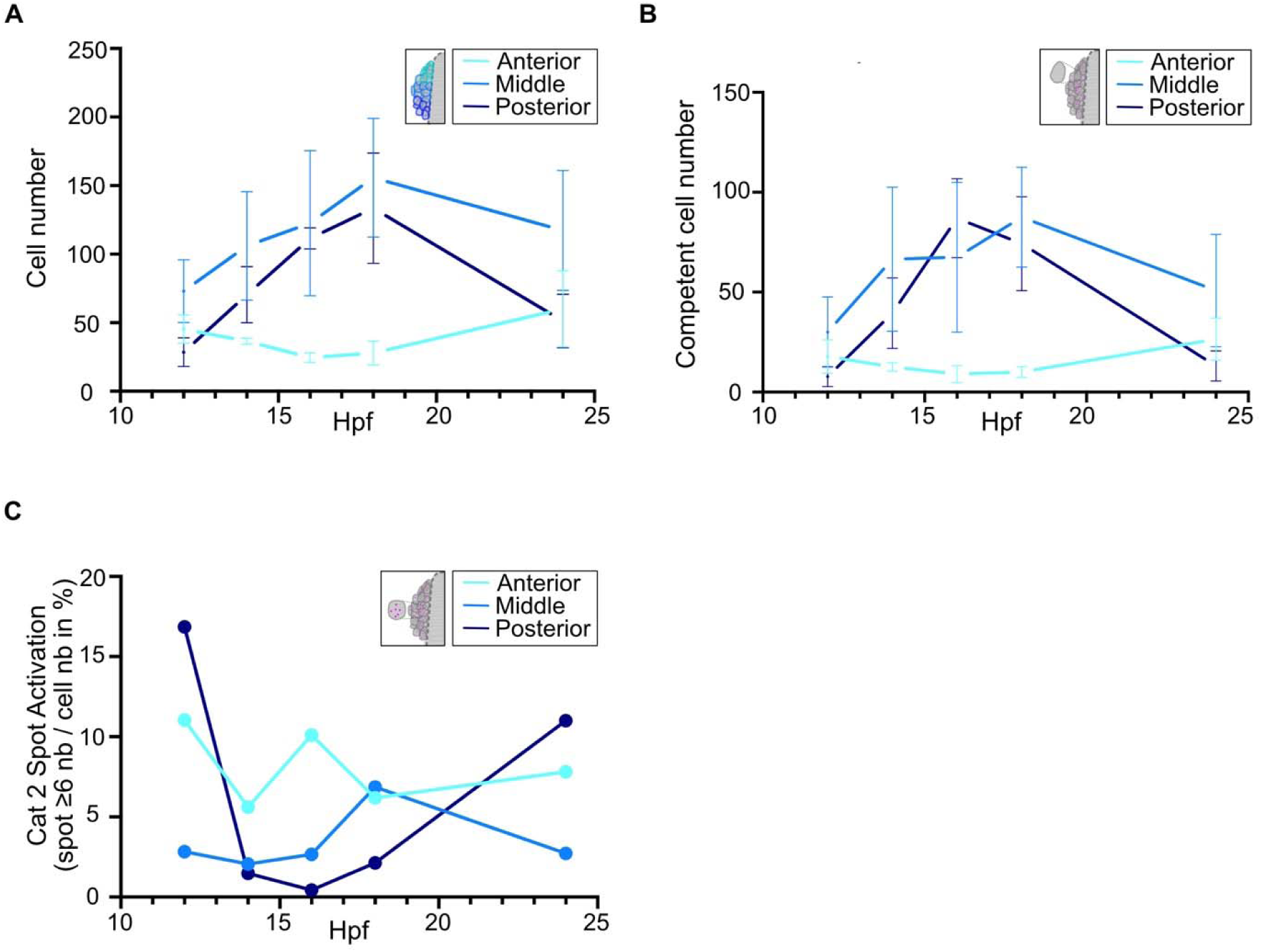
Quantification of the dynamics of Cxcl12a activation during OE formation. (A), Analysis of the number of cells expressing Cxcr4b at the membrane of anterior (light blue), middle (blue) and posterior (dark blue) olfactory cells as a function of time. Shown are mean ±s.d. (B), Representation of the number of cells without any intracellular Cxcr4b+ spot indicating a competent cell as a function of their position along the AP axis. (C), Representation of the number of intracellular Cxcr4b+ category 2 spots as a percentage (*i.e.* number of spots ≥6 per cell, Cat 2 Spot, over the total number of EGFP+ cells) in anterior (light blue), middle (blue) and posterior (dark blue) cells as a function of time.

**Movie S1.** Movie from a simulation in the absence of chemotaxis to mimic the olfactory morphogenesis of the cxcl12a mutant (Fig. 1, D and E, condition 5).

**Movie S2.** Movie of a simulation in the absence of chemotaxis with low Cell-Cell force and low Cell-Matrix force (Fig. 1E, condition 1).

**Movie S3**. Movie of a simulation in the absence of chemotaxis with an intermediate Cell-Cell force and a low Cell-Matrix force (Fig. 1E, condition 2).

**Movie 4.** Movie of a simulation in the absence of chemotaxis with a strong Cell-Cell force and a weak Cell-Matrix force (Fig. 1E, condition 3).

**Movie S5**. Movie of a simulation in the absence of chemotaxis with a low Cell-Cell force and an intermediate Cell-Matrix force (Fig.1E, condition 4).

**Movie S6**. Movie of a simulation in the absence of chemotaxis with a high Cell-Cell force and an intermediate Cell-Matrix force (Fig. 1E, condition 6).

**Movie S7**. Movie of a simulation in the absence of chemotaxis with a low Cell-Cell force and a high Cell-Matrix force (Fig. 1E, condition 7).

**Movie S8**. Movie of a simulation in the absence of chemotaxis with an intermediate Cell-Cell force and a high Cell-Matrix force (Fig. 1E, condition 8).

**Movie S9**. Movie of a simulation in the absence of chemotaxis with a high intermediate Cell-Cell force and a high Cell-Matrix force (Fig. 1E, condition 9).

**Movie S10**. Movie of a simulation in the presence of a source in the whole chemotaxis matrix with intermediate Cell-Cell and Cell-Matrix forces (Fig. 2A, Matrix).

**Movie S11**. Movie of a simulation in the presence of a vertical source with intermediate Cell-Cell and Cell-Matrix forces (Fig. 2D, Vertical).

**Movie S12**. Movie of a simulation in the presence of an horizontal source with intermediate Cell-Cell and Cell-Matrix forces (Fig. 2G, Horizontal).

**Movie S13**. Movie of a simulation in the presence of a central fixed gradient source with intermediate Cell-Cell and Cell-Matrix forces (Fig. 2J, +Chem_Dist).

**Movie S14**. Movie of a simulation in the presence of a central constant force with intermediate Cell-Cell and Cell-Matrix forces (Fig. 2M, +Chem_Const).

**Movie S15**. Movie of a simulation in the presence of an anterior constant force with intermediate Cell-Cell and Cell-Matrix forces (Fig. 4A, Posterior).

**Movie S16**. Movie of a simulation in the presence of a posterior constant force with intermediate Cell-Cell and Cell-Matrix forces (Fig. 4F, Anterior).

